# IL7Rα is required for hematopoietic stem cell reconstitution of tissue-resident lymphoid cells

**DOI:** 10.1101/2021.07.13.452134

**Authors:** Atesh K. Worthington, Taylor S. Cool, Donna M. Poscablo, Adeel Hussaini, Anna E. Beaudin, E. Camilla Forsberg

**Affiliations:** Institute for the Biology of Stem Cells, University of California-Santa Cruz, Santa Cruz, California, United States of America; Program in Biomedical Science and Engineering: Molecular, Cell, and Developmental Biology, University of California-Santa Cruz, Santa Cruz, California, United States of America; Department of Biomolecular Engineering, University of California-Santa Cruz, Santa Cruz, California, United States of America

## Abstract

Traditional, adult-derived lymphocytes that circulate provide adaptive immunity to infection and pathogens. However, subsets of lymphoid cells are also found in non-lymphoid tissues and are called tissue-resident lymphoid cells (TLCs). TLCs encompass a wide array of cell types that span the spectrum of innate-to-adaptive immune function. Unlike traditional lymphocytes that are continuously generated from hematopoietic stem cells (HSCs), many TLCs are of fetal origin and poorly generated from adult HSCs. Here, we sought to understand the development of murine TLCs across multiple tissues and therefore probed the roles of Flk2 and IL7Rα, two cytokine receptors with known roles in traditional lymphopoiesis. Using Flk2- and Il7r-Cre lineage tracing models, we found that peritoneal B1a cells, splenic marginal zone B (MZB) cells, lung ILC2s and regulatory T cells (Tregs) were highly labeled in both models. Despite this high labeling, highly quantitative, in vivo functional approaches showed that the loss of Flk2 minimally affected the generation of these cells *in situ*. In contrast, the loss of IL7Rα, or combined deletion of Flk2 and IL7Rα, dramatically reduced the cell numbers of B1a cells, MZBs, ILC2s, and Tregs both *in situ* and upon transplantation, indicating an intrinsic and more essential role for IL7Rα. Surprisingly, reciprocal transplants of WT HSCs showed that an IL7Rα^-/-^ environment selectively impaired reconstitution of TLCs when compared to TLC numbers *in situ*. Taken together, our data revealed functional roles of Flk2 and IL7Rα in the establishment of tissue-resident lymphoid cells.

## INTRODUCTION

Traditional, circulating immune cells are typically defined as either myeloid or lymphoid and generated from hematopoietic stem cells (HSCs). Myeloid cells are involved in rapid, broad response innate immunity whereas traditional lymphoid cells are involved in slow, specific adaptive immunity [1–4]. The adaptive immune response relies on the complex B and T cell receptor repertoire generation. Although this dichotomy of immune cells is clear for circulating cells, it is unclear if tissue-resident immune cells squarely belong to the myeloid or lymphoid lineage with many recent studies alluding to complex and dynamic origins [4–6]. Unlike circulating immune cells that traffic to non-lymphoid organs upon activation, tissue-resident immune cells reside in non-lymphoid organs, do not recirculate, and have specialized functions that span the innate-adaptive spectrum [7,8]. For example, B1a cells in the peritoneal cavity are a considered B cells [9]. However, they do not undergo the same B cell receptor selection process as circulating B cells and are thus deemed an “innate-like” lymphoid cell. Consequently, it is unclear if TLC differentiation is orchestrated similarly to circulating lymphoid cells; here we sought to determine if their differentiation is regulated by classic lymphoid genes.

We were particularly interested in the cytokine receptors Flk2 and IL7Rα because they are required for traditional adult lymphopoiesis. This finding was demonstrated by impaired lymphopoiesis in both Flk2^-/-^ and Il7rα^-/-^ mice and supported by Flk2-Cre and Il7rα-Cre driven lineage tracing of cells with increasingly restricted lymphoid potential [5,10–12] **(Fig. 1A, 1B, 1C)**. As previously described [5,6,13–15], these models were generated by crossing mice expressing Flk2-Cre [16] or Il7r-Cre [17]to mTmG mice expressing a dual-color fluorescent reporter [18] creating the “FlkSwitch” and “Il7rSwitch” models (**Fig.1A**). In both models, cells express Tomato (Tom) until Cre-mediated recombination results in the irreversible switch to GFP expression by that cell and all of its downstream progeny (**Fig.1B, 1C**). We previously demonstrated that Il7r-Cre labeling is not constrained to lymphoid cells. Surprisingly, in the “Il7rSwitch” lineage tracing model, fetally-derived adult tissue-resident macrophages (TrMacs), were highly labeled and IL7Rα was required for early TrMac development [5], whereas Flk2-Cre did not efficiently label TrMacs [5,19,20]. We also previously examined lung eosinophils, another myeloid cell type; interestingly, despite the high Flk2-Cre and minimal Il7r-Cre labeling of eosinophils, they depend on cell extrinsic IL7Rα, but not Flk2 [15]. The development and generation of these tissue-resident myeloid cells evidently does not follow that of other traditional, circulating myeloid cells. Similarly, many innate-like lymphoid cells are thought to arise from fetal progenitors [4] and it remains unclear if their development follow traditional lymphoid paths and whether they are functionally regulated by known lymphoid drivers. For example, there is evidence that many TLCs arise via common lymphoid progenitor (CLP)-independent pathways [21], and there is differential requirement for Flk2 and IL7Rα amongst different hematopoietic cell types [11,22]. To determine whether Flk2 and IL7Rα are involved in TLC development, we employed lineage tracing, germline knockouts, and HSC transplantation assays.

**Figure 1:**
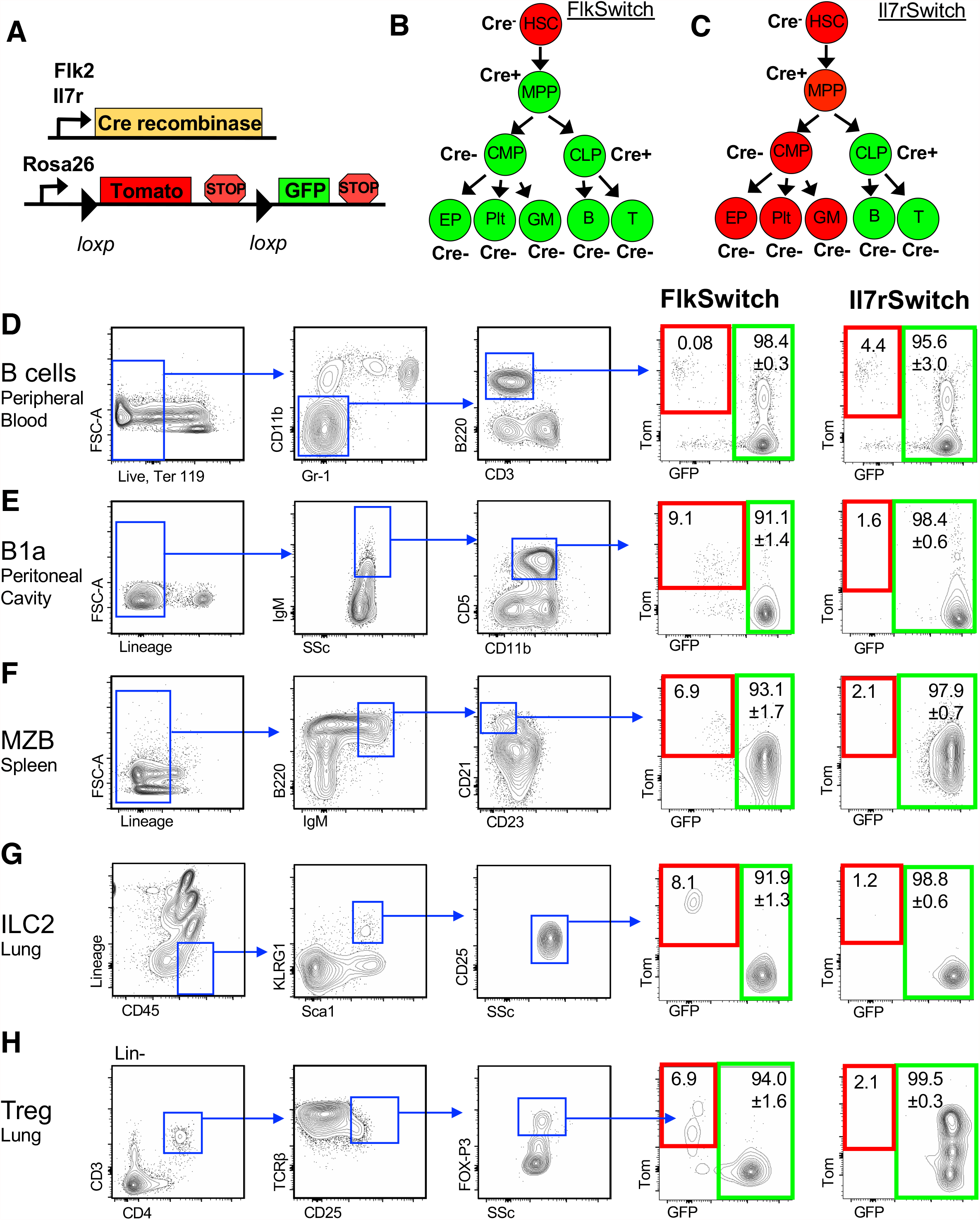
Flk2-Cre & IL7R-Cre efficiently labeled tissue-resident lymphoid populations. **A**) Genetics of the “Switch” models. Flk2 or IL7R regulatory elements drive Cre recombinase expression. Flk2-Cre or IL7r-Cre mice were crossed to Rosa26^mTmG^ mice expressing a dual color reporter expressing either Tomato (Tom) or GFP. **B**,**C)** Schematic of Cre-mediated reporter switching in the “FlkSwitch” (B) and “IL7rSwitch” (C) models. Expression of Cre results in an irreversible switch from Tomato to GFP expression. Once a cell expresses GFP, it can only give rise to GFP-expressing progeny. These models represent Cre-driven labeling in young adult steady state hematopoiesis of circulating, “traditional”, mature myeloid and lymphoid cells. **D-H)** TLCs were highly labeled in both FlkSwitch and IL7rSwitch lineage tracing models. Representative flow cytometric analysis of reporter expression across “traditional” circulating B cells and TLCs, all pre-gated on singlets, lymphocytes and live cells. **D)** “Traditional” B cells (Ter 119^-^ CD11b^-^ Gr-1^-^ CD3^-^ B220^+^) in the peripheral blood and different tissue-resident lymphoid populations in adult mice: **E)** B1a (Lin^-^IgM^+^CD5^+^CD11b^mid^) cells in the peritoneal cavity; **F**) MZB: Marginal Zone B cells (Lin^-^B220^+^IgM^+^CD21^+^CD23^-^) in the spleen; **G)** ILC2: Innate Lymphoid cell Type 2 (Lin^-^CD45^+^KLRG^+^Sca-1^+^CD25^+^) in the lung; and **H)** Tregs: regulatory T cells (Lin^-^CD3^+^CD4^+^TCRB^+^CD25^+^FOX-P3^+^) in the lung. Tom and GFP expression are highlighted by red and green boxes, respectively, in FlkSwitch and IL7RSwitch models. Values indicate mean frequencies ± SEM of gated Flk2-Cre and IL7R-Cre marked GFP+ populations. Data are representative of 4-5 mice per cohort in three independent experiments.

## MATERIALS & METHODS

### Mice

All animals were housed and bred in the AAALAC accredited vivarium at UC Santa Cruz and group housed in ventilated cages on a standard 12:12 light cycle. All procedures were approved by the UCSC Institutional Animal Care and Use (IACUC) committees (OLAW assurance A3859-01; USDA Registration 93-R-0439, customer number 9198). Il7r-Cre [17], and Flk2-Cre [16] mice, obtained under fully executed Material Transfer Agreements, were crossed to homozygous Rosa26^mTmG^ females (JAX Stock # 007576) [18] to generate “switch” lines, all on the C57Bl/6 background. WT C56Bl/6, Rosa26^mTmG^ and UBC-GFP (JAX Stock # 004353) [23] mice were used for controls. FIDKO (**F**lk2 **I**l7rα **D**ouble **K**nock-**O**ut) mice were generated by crossing Flk2^-/-^ [24] and IL7Rα ^-/-^ [12] mice to homozygosity. Adult male and female mice (8-12 weeks old) were used randomly and indiscriminately, with the exception of the FlkSwitch line, in which only males were used because many female mice do not carry a Cre allele.

### Tissue and cell isolation

Mice were sacrificed by CO2 inhalation. Adult lung and spleen were isolated, weighed, then placed into 1.5mL of digestion buffer (1x PBS(+/+) with 2% serum, 1 mg/mL (lung) or 2mg/ml (spleen) collagenase IV (Gibco) with 100U/ml DNase1) containing Calibrite APC-labeled (BD Biosciences) counting beads and manually disassociated into 2mm x 2mm pieces using surgical scissors. The tissues were then incubated at 37°C for 1 hour (lung) or 2 hours (spleen). Following incubation, all tissues were passaged through 19g then a 16g needle 10 times, and then filtered through a 70 µM filter and 25mL of cell staining buffer (1X PBS with 2% serum and 5mM EDTA) was added to quench the digestion enzymes.

### Flow Cytometry & Cell Analysis

Cell labeling was performed on ice in 1X PBS with 5 mM EDTA and 2% serum. Analysis was performed on a BD FACS Aria III (BD Bioscience, San Jose, CA) at University of California-Santa Cruz, and analyzed using FlowJo (Treestar)[25–27]. Cells were defined as follows: B1a = Lin^-^ (Ter119, CD11b, CD3) IgM^+^ CD5^+^ CD11b^mid^; MZB = Lin^-^(CD3, CD4, CD8, Ter-119, Gr1) B220^+^ IgM^+^ CD21^+^ CD23^-^; ILC2 = Lin^-^ (CD3, CD4, CD5, CD8, NK1-1, CD19, Ter119, F4/80, FcεRIα) CD45^+^ KLRG1^+^ Sca-1^+^ CD25^+^; Treg = Lin^-^ (Ter119, Gr1) CD3^+^ CD4^+^ TCRB^+^ CD25^+^ FOX-P3^+^.

### Transplantation Assays

Transplantation assays were performed as previously described [6,11,28,29]. Briefly, double sorted HSCs were isolated from bone marrow of either WT (non-fluorescent, mTmG or UBC-GFP), Flk2^-/-^, Il7rα^-/-^ or FIDKO 8-12 week old mice. HSCs were defined as KLS (cKit^+^Lin^-^Sca-1^+^), SLAM^hi^Flk2^-^. Lineage markers were CD3, CD4, CD5, CD8, B220, Mac-1, Gr-1 and Ter-119. Flk2 could not be used to identify HSCs in Flk2^-/-^ and FIDKO HSCs and so they were sorted on KLS, SLAM^hi^. Recipient mice aged 8-12 weeks were sublethally irradiated (750 rad, single dose) with a Faxitron CP-160 (Faxitron). Under isofluorane-induced general anesthesia, sorted cells were transplanted retro-orbitally. Recipient mice were bled 4, 8, 12, and 16 weeks post transplantation via tail vein and peripheral blood (PB) was analyzed for donor chimerism by means of fluorescence profiles and antibodies to lineage markers (Table 1). Long-term multilineage reconstitution was defined as chimerism in both the lymphoid and myeloid lineages of > 0.1% at 16 weeks post-transplantation and only mice that displayed long-term reconstitution were used for post-transplantation analysis.

**TABLE 1.** Antibodies used in experiments.

| Antibody | Source | Identifier | Antibody | Source | Identifier |
| --- | --- | --- | --- | --- | --- |
| B220 - APC Cy7 | Biolegend | 103224 | CD3 - A700 | Biolegend | 100216 |
| CD3- APC | Biolegend | 100236 | CD4 - A700 | Biolegend | 100429 |
| CD4 - BV605 | Biolegend | 100451 | CD5 - A700 | R&D Systems | FAB115N-025 |
| CD5 - APC | Biolegend | 100625 | CD8 - A700 | Biolegend | 300919 |
| CD11b - PB | Biolegend | 101223 | Ter119 - A700 | Biolegend | 116220 |
| CD11b- PECy7 | Biolegend | 101216 | B220 - A700 | Biolegend | 103231 |
| CD21 - PE Cy7 | Ebiosciences | 25-0211-82 | Gr1 - A700 | Biolegend | 108421 |
| CD23 - PB | Biolegend | 101616 | CD11b - A700 | Biolegend | 101222 |
| CD25 - APC Cy7 | Biolegend | 102026 | CD3 - biotin | Biolegend | 100243 |
| CD45.2 - pB | Biolegend | 109820 | CD4 - biotin | Biolegend | 100403 |
| CD61-Alexa 647 | Biolegend | 104314 | CD5 - biotin | Biolegend | 100603 |
| CD150 - BV786 | Biolegend | 115937 | CD8 - biotin | Biolegend | 100703 |
| cKit - APC Cy7 | Biolegend | 105826 | Ter119 - biotin | Biolegend | 116203 |
| Flk2 - APC | Biolegend | 135310 | Nk1.1 - biotin | Biolegend | 108703 |
| FoxP3 - PB | Biolegend | 126409 | CD19 - biotin | Biolegend | 115503 |
| Gr1-Pacific Blue | Biolegend | 108430 | F4/80 - biotin | Biolegend | 123105 |
| IgM - BV605 | Biolegend | 406523 | FcεRIα - biotin | Biolegend | 101303 |
| KLRG - BV605 | Biolegend | 138419 | STA - BV786 | Biolegend | 405249 |
| Sca1 - PB | Biolegend | 122520 |  |  |  |
| TCRB - Pe Cy7 | Biolegend | 109222 |  |  |  |
| Ter119-PECy5 | Biolegend | 116210 |  |  |  |

### Absolute Cell Number Quantification

A known volume of PB was mixed with an antibody solution, (PBS, 5mM EDTA and 2% serum) containing a known quantity of Calibrite APC beads prior to flow cytometry analysis [15,25,30]. For tissues, a known quantity of beads was added to each at the very beginning of tissue preparation prior to antibody staining and analysis. The number of beads counted by flow cytometry was used to calculate the number of mature cells per micro liter of blood or within each tissue.

### Quantification and Statistical Analysis

Number of experiments, n, and what n represents can be found in the legend for each figure. Statistical significance was determined by two-tailed unpaired student’s T-test. All data are shown as mean ± standard error of the mean (SEM) representing at least three independent experiments.

## RESULTS

### Flk2-Cre and Il7r-Cre highly label tissue-resident lymphoid cell populations

To test whether TLCs arise via differentiation pathways similar to circulating lymphocytes, we examined labeling of TLC labeling the FlkSwitch and Il7rSwitch lineage tracing models (**Fig. 1A-C**). We compared reporter expression of traditional peripheral blood B cells to reporter expression of B1a cells isolated from the peritoneal cavity, marginal zone B cells (MZB) from the spleen, type 2 innate lymphoid cells (ILC2) from the lung, and regulatory T cells (Treg) from the lung (**Fig. 1D-H**). As expected, and previously reported, Cre-driven labeling of circulating peripheral blood B cells was greater than 95% in both FlkSwitch and Il7rSwitch mice (**Fig. 1D**) [5,17]. Similarly, Cre-driven labeling was uniformly high for all TLCs examined, a trend we also observed in B1b and B2 cells (**Fig. S1A**). Together, our data showed that B1a cells, ILC2, MZB and Tregs arise from Flk2- and IL7Rα-positive cells, as do traditional circulating lymphoid cells.

### Tissue-resident lymphoid cells are severely reduced in the absence of IL7Rα, but not Flk2

We and others have previously shown that loss of Flk2 results in a reduction of hematopoietic progenitor cells, with a less severe reduction in mature B cells in the peripheral blood [11,22,24,31]. The loss of IL7Rα has also been shown to result in a severe reduction of peripheral blood B cells [12]. Here, we observed similar trends in the lung, where traditional CD19+ B cells, although not significantly reduced in Flk2^-/-^ mice, were significantly reduced in the IL7Rα^-/-^ mice (**Figure 2B**). To determine if TLCs were similarly affected, we quantified their cellularity in the absence of Flk2 and IL7Rα. Compared to WT numbers of each cell type, only MZBs were significantly reduced in the absence of Flk2 (**Fig. 2D**), whereas B1a cells, ILC2s and MZBs were all significantly reduced in the absence of IL7Rα (**Fig. 2C, 2D, 2E; Fig. S2A**), consistent with prior studies of these cells in the same and other tissues, and peritoneal B1b and B2 cells (**Fig. 1SB-B’**) [32–35]. Interestingly, despite being as efficiently labeled by both Flk2-Cre and IL7r-Cre (**Figure 1H**), Tregs were not significantly reduced in either Flk2^-/-^ or IL7Rα^-/-^ mice, although there was a downward trend (**Fig. 2 F**), as previously seen in thymic and splenic Tregs of IL7Rα^-/-^ mice [36]. We hypothesized that Flk2 may compensate for the lack of IL7Rα, and vice versa, allowing for near-normal Treg development in the absence of either receptor. To test this hypothesis, we generated **<u>F</u>**lk2, **<u>I</u>**L7Rα **<u>d</u>**ouble **<u>k</u>**nock**<u>o</u>**ut (FIDKO) mice and quantified total Tregs in the lung. We found a significant reduction of Tregs in the lung of FIDKO mice compared to WT mice (**Fig. 2F**), revealing an overlapping and cell-specific role of Flk2 and IL7Rα. We also quantified B1as, ILC2s and MZBs in the FIDKO mice and found that they recapitulated the severe reductions in numbers that we found in the IL7Rα^-/-^ mice. These data suggest that both Flk2 and IL7Rα do have functional roles in TLC development during steady state hematopoiesis, although IL7Rα appears to be more important than Flk2 as indicated by the more severe reduction in cell numbers.

**Figure 2:**
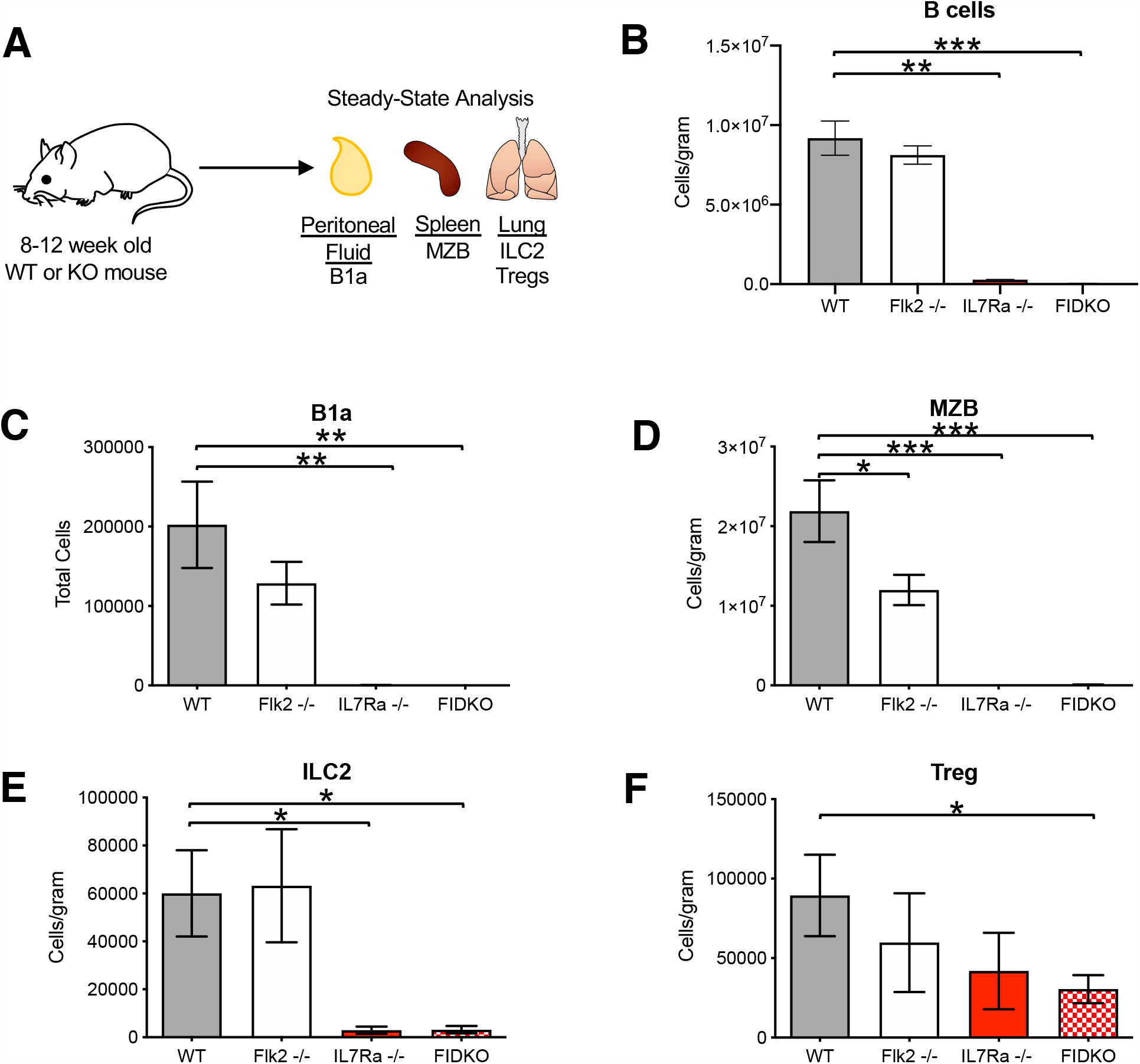
Tissue-resident lymphoid cells are severely reduced in the absence of IL7Rα, but not Flk2. **A)** Schematic of experimental design. Peritoneal fluid, spleen and lung from 8 to 12 week old WT, Flk2^-/-^, IL7Rα^-/-^ and FIDKO HSCs were harvested and analyzed for cellularity of TLCs and represented in bar plots of. WT (grey), Flk2^-/-^ (white), IL7Rα^-/-^ (red) and Flk2^-/-^/IL7Rα^-/-^ double knockout (FIDKO; red/white). **B)** “Traditional” B cells in the lung were significantly reduced in both IL7Rα^-/-^ and FIDKO mice, but not in Flk2^-/-^ mice. Quantification of B cells (Live, Ter119-Mac1-Gr1-CD19+) per gram of tissue in the lungs. **C)** B1a cells in the peritoneal cavity were significantly reduced in IL7Rα^-/-^ and FIDKO mice compared to WT. Quantification of total cell numbers in the peritoneal cavity. **D)** MZBs were significantly reduced in Flk2^-/-^, IL7Rα^-/-^ and FIDKO compared to WT mice. Quantification of cells/gram of tissue in the spleen. **E)** ILC2s were significantly reduced in IL7Rα^-/-^ and FIDKO compared to WT mice. Quantification of cells/gram of tissue in the lung. **F)** Tregs were significantly reduced only in FIDKO compared to WT mice. Quantification of cells/gram of tissue in the lung. Minimum of n=4 per genotype, representing four independent experiments. Differences were analyzed with two-tailed Student t test *, P<0.05; **, P< 0.005; ***, P< 0.0005

### Flk2^-/-^ and IL7Rα^-/-^ HSCs have impaired tissue-resident lymphoid cell reconstitution

Hematopoietic stem and progenitor cell (HSPC) differentiation relies on cytokine receptors such as Flk2 and IL7R to receive instructive signals from the environment [37]. For example, we have previously shown that Flk2 is intrinsically required on HSPCs for both myeloid and lymphoid reconstitution, as demonstrated by the reduced capacity of Flk2^-/-^ HSCs to generate myeloid and lymphoid progeny, including traditional circulating lymphoid cells, upon transplantation into a WT host [11]. More recently, we demonstrated an extrinsic requirement of IL7Rα in the generation of eosinophils from WT HSCs, as demonstrated by a reduced capacity of WT HSCs to generate eosinophils upon transplantation into an IL7Rα^-/-^ host [15]. Therefore, we were curious if the deficiency in TLCs we observed in Flk2^-/-^, IL7Rα^-/-^ and FIDKO mice (**Figure 2**) is due to the cell-intrinsic lack of receptors or due to changes to surrounding cells. To begin to answer this question, we transplanted WT, Flk2^-/-^, IL7Rα^-/-^ or FIDKO HSCs into irradiated wildtype hosts (**Fig. 3A**). WT HSCs are Lin^-^cKit^+^Sca^+^SLAM^+^Flk2^-^, therefore we were able to isolate HSCs from all mice with the same markers (**Figure S3**). We first examined donor chimerism of circulating lymphoid cells. As expected, peripheral blood B cell donor chimerism was significantly lower from Flk2^-/-^ donor HSCs, IL7Rα^-/-^ HSCs and FIDKO HSCs compared to WT HSCs (**Fig. 3B**). When examining TLCs, we observed only donor chimerism of from IL7Rα^-/-^ and FIDKO HSCs was significantly impaired, while Flk2 deficiency alone did not result in significant differences (**Fig. 3C-D, Fig. S1C**,**C’’, Fig. S2B**). However, a limitation of donor chimerism is that this measure relies on host cell numbers and therefore makes it more difficult to determine the intrinsic requirement of these receptors if the number of host cells change [25,30]. To overcome this limitation, we quantified the absolute cellularity of donor-derived cells, which is purely a measure of donor cells. Quantification of absolute cellularity of donor-derived cells between WT and knockout HSCs [25,30] revealed that reconstitution of B1a cells (**Fig. 3C’**) and MZBs (**Fig. 3D’**) was also significantly impaired by the loss of Flk2 alone. Regardless of quantification method, deletion of either IL7Rα alone, or both IL7Rα and Flk2, led to impaired reconstitution of all TLCs examined (**Fig. 3 C’-F’; Fig. 2SB’**). This was intriguing since we did not observe significant reduction of Tregs in IL-7Rα^-/-^ at steady state (**Fig. 2F**). This is likely due the stress of transplantation which reveals more dynamic requirement of IL-7Rα, as we previously observed a similar trend when we compared granulocytes/macrophages at steady state in Flk2^-/-^ mice to transplantation of Flk2^-/-^ HSCs [11]. These data suggest that IL7Rα is required cell intrinsically for TLC reconstitution, that Flk2 cannot compensate for this requirement, and that IL7Rα is not capable of fully compensating for the loss of Flk2.

**Figure 3:**
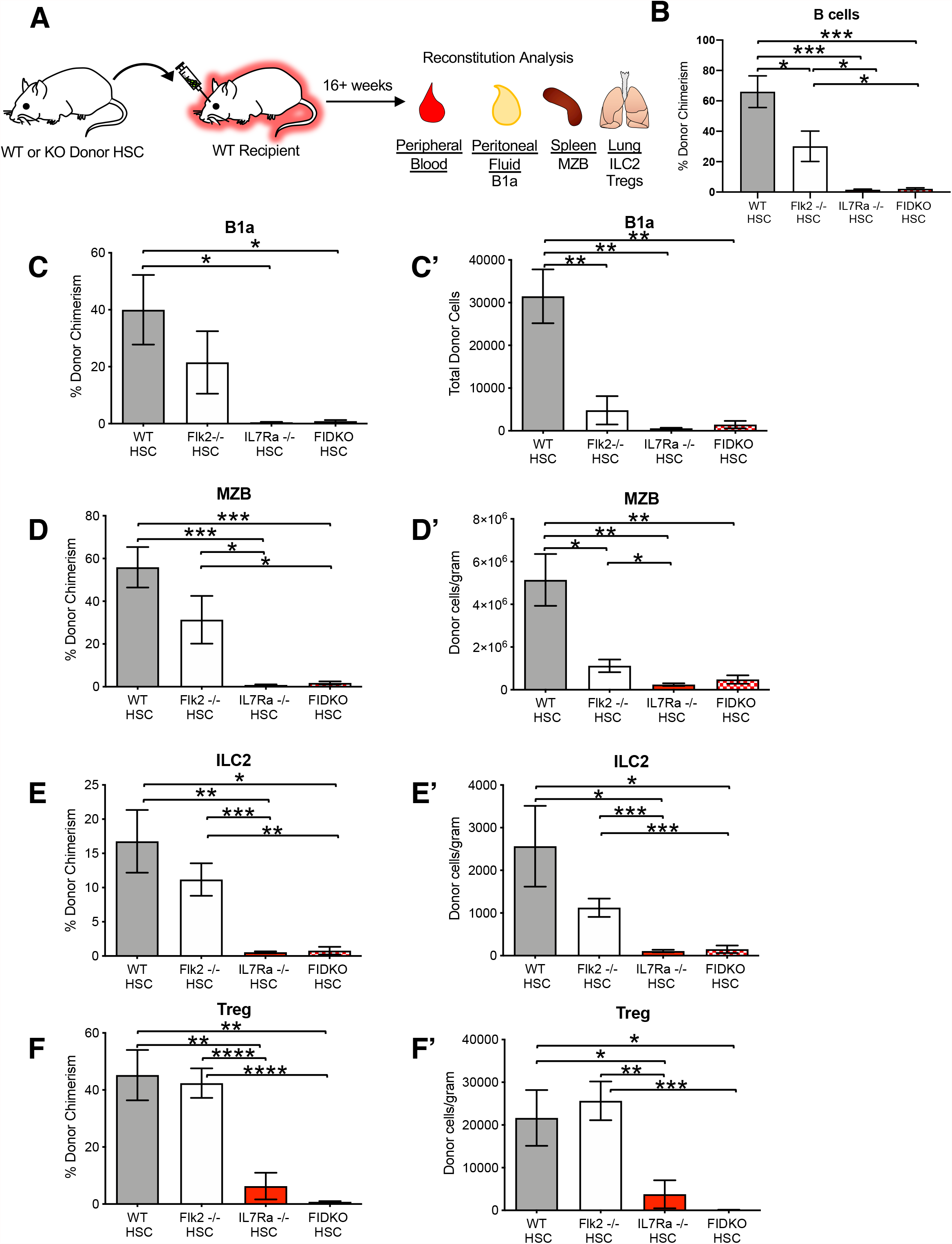
IL7Rα^-/-^ and Flk2^-/-^ HSCs had impaired tissue-resident lymphoid cell reconstitution compared to WT HSCs. **A)** Schematic of experimental design. 500 WT, Flk2^-/-^, IL7Rα^-/-^ or FIDKO HSCs were transplanted into sublethally irradiated fluorescent WT recipients (mTmG or UBC-GFP) and donor contribution of TLCs was quantified 16 weeks post transplantation. **B)** Reconstitution of peripheral blood B cells was significantly impaired upon transplantation of Flk2^-/-^(white bars), IL7Rα^-/-^ (red bars), or FIDKO HSCs (red/white bars) compared to WT HSCs (grey bars). Quantification of donor chimerism of “traditional” B cells in the peripheral blood. **C-F)** Donor chimerism of TLCs significantly reduced from IL7Rα^-/-^ (red bars) and FIDKO (red square bars) donor HSCs compared to WT (grey bars) donor HSCs. Quantification of donor chimerism of B1a cells, MZBs, ILC2s and Tregs, respectively. **C’-F’)** Cellularity of B1a and MZB cells were significantly reduced from Flk2^-/-^ (white bars), IL7Rα^-/-^ (red bars) and FIDKO (red square bars) donor HSCs compared to WT (grey bars) donor HSCs. Cellularity of lung ILC2 and Tregs were significantly reduced from, IL7Rα^-/-^ (red bars) and FIDKO (red square bars), but not Flk2^-/-^ (white bars), donor HSCs compared to WT (grey bars) donor HSCs. Quantification of total B1a cells, cells/gram spleen of MZBs, cells/gram lung ILC2s and cells/gram of lung tissue Tregs, respectively. WT n=9, Flk2^-/-^ n=11, IL7Rα^-/-^ n=7, FIDKO n=4, representing a minimum of four independent experiments. Differences were analyzed with two-tailed Student t test *, P< 0.05; **, P< 0.005; ***, P< 0.0005.

### WT HSCs have enhanced tissue-resident lymphoid cell reconstitution capacity in an IL7Rα^-/-^ environment

Although the HSC transplantations of Figure 3 demonstrated a cell intrinsic requirement of IL7Rα for TLC reconstitution, it is also possible that extrinsic IL7Rα may be required, as we previously demonstrated for eosinophil reconstitution [15]. To test this, we performed transplantation of WT HSCs into IL7Rα-deficient mice (**Fig. 4A**). Surprisingly, we observed significantly greater numbers of donor-derived B1a cells, MZBs, ILC2s and Tregs (**Fig. 4B-E, Fig. S2C)**, as well as B1b and B2 cells (**Fig. S1D, D’’**), in the IL7Rα^-/-^ hosts compared to WT hosts. This raised the question of whether the greater number of donor-derived cells in the IL7Rα^-/-^ mice was reflective of an overall greater number of cells, or if the reconstitution by WT HSCs only compensated for the lower cell numbers of cells in the IL7Rα^-/-^ host. Therefore, we determined the total number of host and donor derived cells and compared these numbers between WT hosts and IL7Rα^-/-^ hosts. We found no significant differences between WT and IL7Rα^-/-^ recipients of total B1a (**Fig. 4B’**) and MZB (**Fig. 4C’**) cells, or B1b and B2 cells (**Fig. S1D’**,**D’’’**). Interestingly, we observed significantly greater total numbers of ILC2s (**Fig. 4D’, Fig. 2SC’**) and Tregs (**Fig. 4E’**) in the IL7Rα^-/-^ recipients compared to the WT hosts. The enhanced reconstitution of tissue-resident T lymphocytes in an IL7Rα^-/-^ environment compared to consistent reconstitution of tissue-resident B lymphocytes suggests differential dependence on IL-7/IL7R signaling in the development of these cell types [38–40].

**Figure 4:**
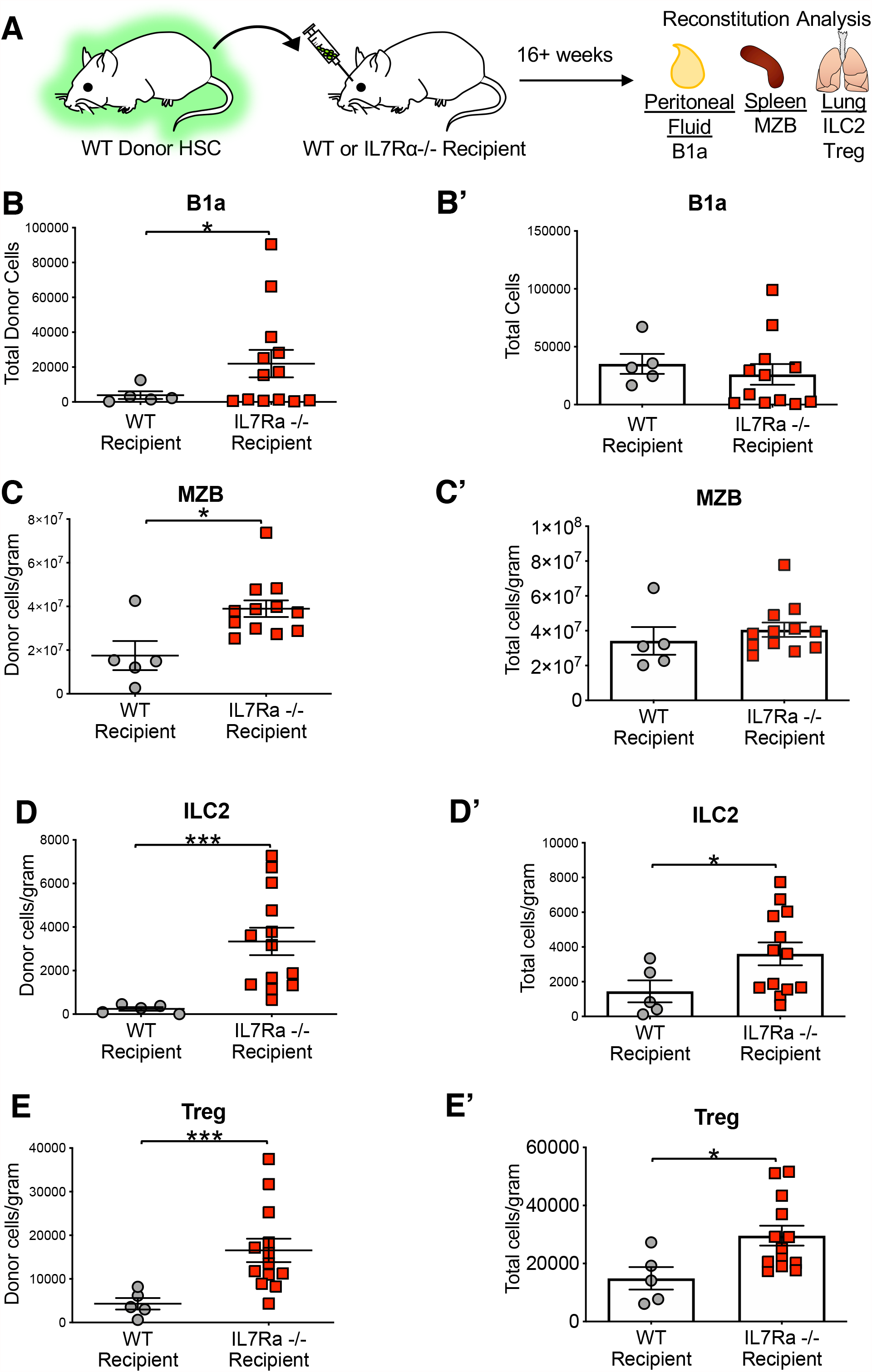
WT HSCs have enhanced non-traditional lymphoid cell reconstitution capacity in an IL7Rα^-/-^ environment. **A)** Schematic of experimental design. 500 fluorescent WT HSCs (UBC-GFP), were transplanted into sublethally irradiated non-fluorescent WT or IL7Rα^-/-^ recipients. Donor contribution to TLCs was quantified 16 weeks post-transplantation. **B-E)** Significantly greater donor-derived TLCs in an IL7Rα^-/-^ recipient (red squared) compared to a WT recipient (grey dots). Quantification of donor-derived cells of B1a cells, ILC2s, MZBs and Tregs, respectively. **B’-E’)** Significantly greater total ILC2 and Treg cells in an IL7Rα^-/-^ recipient (red squared) compared to a WT recipient (grey dots). Quantification of total cells (donor+host) of B1a cells, ILC2s, MZBs and Tregs, respectively. WT recipient n=5, IL7Rα^-/-^ recipient n=13, representing four independent experiments. Differences were analyzed with two-tailed Student t test *, P<0.05; **, P< 0.005; ***, P< 0.0005.

### WT HSCs transplanted into IL7Rα^-/-^ mice are not fully capable of rescuing the impaired lymphoid phenotype

The majority of TLCs are thought to arise pre/perinatally, with little contribution from adult progenitors. Although we observed enhanced reconstitution of some TLCs by adult WT HSCs in IL7Rα^-/-^ hosts, it remained unclear if the lymphoid deficiencies observed in the IL7Rα^-/-^ mice (**Fig. 2**) were fully rescued by transplantation of adult WT HSCs. Therefore, we compared steady-state numbers of circulating and TLCs in both WT and IL7Rα^-/-^ mice (**Fig. 2**, solid bars) to total numbers after long-term hematopoietic reconstitution post-transplantation of WT HSCs (**Fig. 4B’-4E’)**. In WT and IL7Rα^-/-^ mice transplanted with WT HSCs, we found no significant difference in total peripheral blood B cells compared to WT mice at steady state (**Fig 5A**), as well in B1b and B2 cells (**Fig.1SE-E’**). Similarly, we observed no difference between total numbers of MZBs in WT mice at steady-state and WT mice transplanted with WT HSCs (**Fig. 5C**, solid grey bar compared to grey lined bar). Therefore, WT HSCs were indeed capable of fully reconstituting these cells. In fact, although the difference in total MZBs was not significantly different between WT and IL7Rα^-/-^ recipients (**Fig. 4C’**), we observed significantly more MZBs in the IL7Rα^-/-^ recipient mice compared to WT steady-state (**Fig. 5C**, solid grey bar compared to red lined bar). Interestingly, this was not the case for the other cell types examined. We observed significantly fewer B1a cells, ILC2s and Tregs in mice transplanted with WT HSCs compared to steady state WT mice (**Fig 5B, 5E-E**; solid grey bars compared to grey and grey striped bars), consistent with reports by us and others of limited TLC potential of adult progenitors [6,41,42]. Despite the enhanced reconstitution of ILC2 and Tregs observed in the IL7Rα^-/-^ environment (**Fig 4 D’-E’**), we were surprised to find that this was not sufficient to return cell numbers to those observed in WT steady-state mice; rather, the total number of cells was not greater than the total numbers observed at steady-state in the IL7Rα^-/-^ mice (**Fig. 5E** & **5E**, solid red bars compared to red lined bars). Taken together, these data suggest that adult WT HSCs transplanted into IL7Rα^-/-^ mice are not fully capable of rescuing the impaired lymphoid phenotype caused by IL7Rα deletion and that there is differential requirement for IL7Rα on the development of tissue-resident B and T cells.

**Figure 5:**
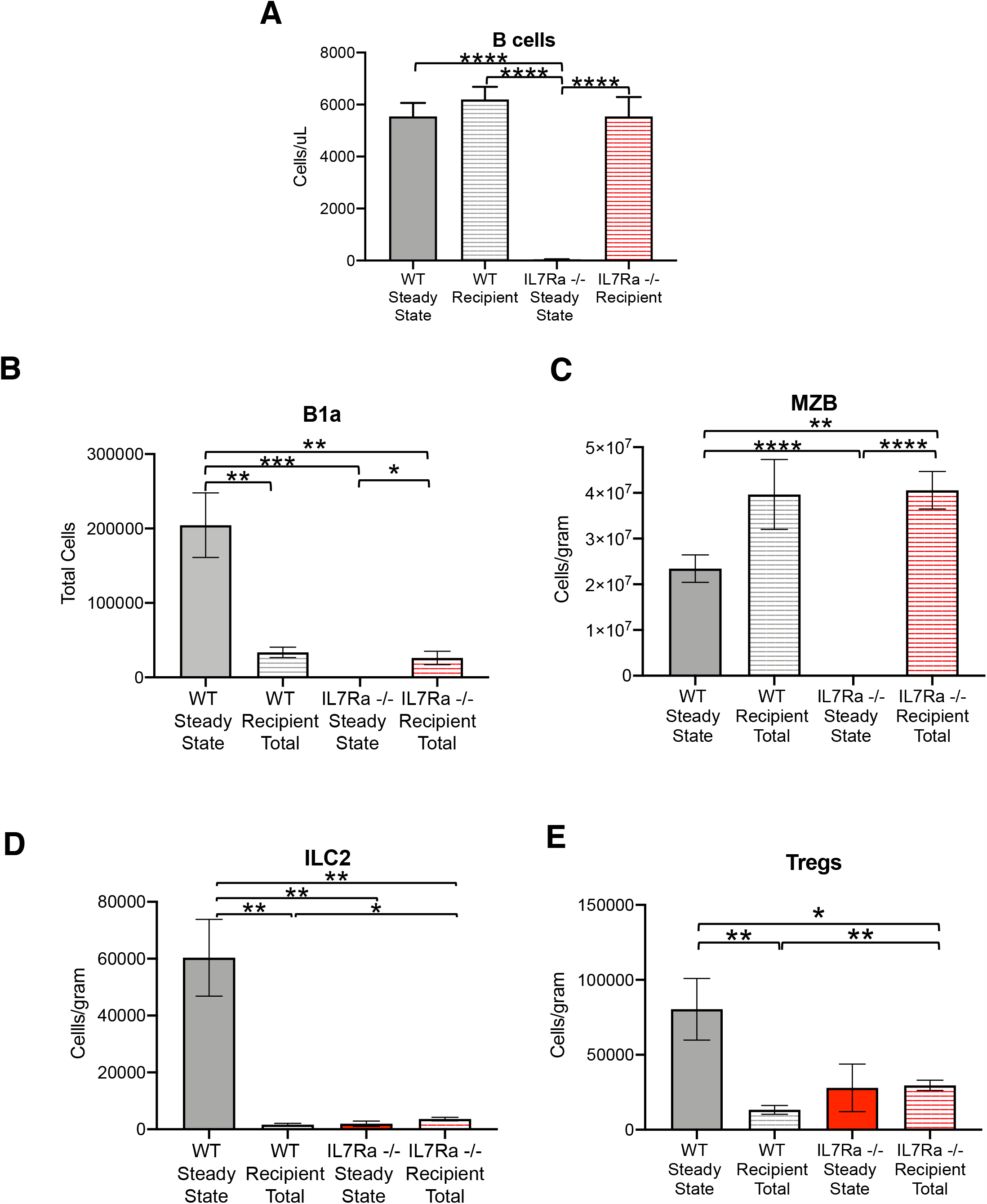
WT HSCs transplanted into IL7Rα^-/-^ mice are not capable of reconstituting tissue-resident lymphocytes to steady state numbers (or levels). Bar graph comparing circulating B cells and TLCs cells at steady state (solid bars) to their reconstitution upon transplantation of WT HSCs (pattern bars). **A)** Cellularity of peripheral blood B cells, **B)** B1a cells, **C)** MZB, **D)** ILC2 and **E)** Tregs in WT or IL7Rα^-/-^ mice are compared to WT and IL7Rα^-/-^ mice reconstituted with WT HSCs. WT Steady state n=11; WT Recipient Total n=7; IL7Rα^-/-^ steady state n=6; IL7Rα^-/-^ Recipient Total n=12. Differences were analyzed with One-way ANOVA; PB B cells p<0.0001; B1a cells p<0.0001; MZB p<0.0001; ILC2 p<0.0001; Tregs = 0.0134. Dunnett multi-parameter test *, P<0.05; **, P< 0.005; ***, P< 0.0005; ****, P< 0.00005.

## DISCUSSION

### Cell-intrinsic IL7Rα is required for differentiation of tissue-resident lymphoid cells

In this study, we aimed to expand our understanding on the roles of the cytokine receptors Flk2 and IL7Rα beyond traditional, circulating lymphocytes by investigating their roles in TLC development and reconstitution. Using lineage tracing, knockout mouse models and transplantation assays, our data demonstrated a clear role of IL7Rα in the differentiation of tissue-resident lymphoid cells across multiple tissues. B1a cells, MZBs, ILC2 and Tregs all displayed very high labeling in both the FlkSwitch and Il7rSwitch lineage tracing models; however, only IL7Rα^-/-^ resulted in severely impaired TLC development while the effects of Flk2^-/-^ were minimal. Furthermore, IL7Rα^-/-^ HSCs were incapable of reconstituting TLCs in WT hosts, while WTs were capable of TLC reconstitution in IL7Rα^-/-^ hosts, revealing a primarily cell-intrinsic requirement of IL7Rα for TLC differentiation.

### Flk2 is less essential for tissue-resident lymphocytes than circulating lymphocytes

Based on the high labeling in the Flk2Switch model (**Fig. 1**) and our previous finding that multipotent, myeloid and lymphoid progenitors, and PB B cells were reduced in Flk2^-/-^ mice [11], we expected to find significantly reduced numbers of TLCs in Flk2-deficient mice. We were surprised that only MZBs were significantly reduced in the Flk2^-/-^ compared to WT mice (**Fig. 2**). We have also previously shown that multilineage reconstitution is impaired upon transplantation of Flk2^-/-^ HSCs in WT recipients, likely due to the significant decrease in HSC differentiation into Flk2+ MPPs and CLPs [11]. Despite demonstrating that B1a cells, MZBs, ILC2 and Tregs differentiate via a Flk2 positive stage (**Fig. 1**), we were surprised that only B1a and MZBs reconstitution was impaired by the loss of Flk2 (**Fig. 3**). This may be due to a more stringent reliance on Flk2 for innate-like B cell rather than innate-like T cells, or reported CLP-independent differentiation [21]. However, previous studies have shown that Tregs increase in number in response to Flt3 ligand in mice and humans [43,44], making it all the more surprising that Tregs are the least affected by the loss of Flk2. These data suggest that IL7Rα is able to compensate for the loss of Flk2 in a cell-type specific manner but not vice versa, and therefore plays a more essential role in TLC reconstitution.

### Absolute cell quantification revealed unexpected reconstitution capability

In all transplantation experiments, we employed an absolute cell quantification method previously developed in the lab [5,15,25]. This method was critical to our interpretation of WT HSC reconstitution capability in IL7Rα^-/-^ recipients because it cannot be gleaned from percent donor chimerism alone. Since the IL7Rα^-/-^ recipients were already devoid of any of the cells we were examining, donor-derived cell would be constituting 100% donor chimerism. By quantifying absolute cell numbers, we found that not only did WT HSCs generate TLCs in an IL7Rα^-/-^ recipient, but that the overall numbers of ILC2 and Tregs were greater in the IL7Rα^-/-^ recipients (**Fig. 4**). We suspect this is due to the abundance of IL-7 in the IL7Rα^-/-^ mice [15,45] and lack of host competition for IL-7, which may in turn enhance the ILC2 and Treg reconstitution. Therefore, IL7Rα may have a cell non-autonomous role in regulating TLC subsets which rely more heavily on IL-7 signaling by controlling IL-7 levels [15,45]. Additionally, in our previous examination of eosinophil reconstitution in IL7Rα^-/-^ mice, we discovered that the concentrations of other cytokines involved in immune cell survival are different in IL7Rα^-/-^ compared to WT mice [15]. Therefore, we speculate the differences in cytokine signals may play a significant role in the differences we observed in Figure 4.

We were also able to compare total TLCs reconstituted by WT HSCs to their total numbers at *in situ* at steady-state. With the exception of MZBs in the IL7Rα ^-/-^ host, WT HSCs in either the WT host or the IL7Rα ^-/-^ host did not fully rescue TLCs to WT steady state levels (**Fig. 5**). This is not altogether very surprising considering we performed our transplants with adult HSCs and that at least some types of TLCs are thought to originate from fetal/neonatal progenitors. For example, we have previously shown that the reconstitution capacity of B-1a cells in the peritoneal cavity and MZBs is significantly greater in a pool of transient, developmentally restricted HSCs compared to adult HSCs [6]. However, it has remained controversial to what extent adult HSCs contribute to B1b cells. Here, we see that B1b cell reconstitution mirrors peripheral blood B cells, indicating that adult HSCs generate them similarly (**Fig. 1SE, Fig. 5A**). Additionally, recent work from Locksley and colleagues has shown that ILC2s from the lung originate from perinatal progenitors, which is also supported by our data (Fig. 5D) [41]. However, the mechanisms that control TLC potential in adult HSCs is poorly understood and deserves further investigation, considering that HSCs are used in the clinic to treat a multitude of immune deficiencies.

### IL7Rα is required for both innate and adaptive immunity

Despite some TLCs displaying immune properties reminiscent of myeloid cells [46–50], we demonstrated here that they nevertheless rely on main regulators of traditional, circulating lymphoid cells. Interestingly, IL7Rα regulation of TLC generation is predominantly cell intrinsic, similar to that of fetal-specified tissue-resident macrophages [5] but in contrast to the solely cell extrinsic role we discovered for IL7Rα in adult myeloid eosinophil regulation [15]. Collectively, these data indicate broad and temporally dynamic functional roles of both Flk2 and IL7Rα beyond the typical lymphoid ascribed roles.

## Acknowledgments

We thank Dr. I. Lemischka for the Flk2^-/-^ mice; Drs. H-R. Rodewald and S.M. Schlenner for the IL7Rα-Cre strain; Dr. T. Boehm for the Flt3^Cre^ strain; Bari Nazario and the UCSC Institute for the Biology of Stem Cells for flow cytometry support.

## Author contributions

A.W., and E.C.F. conceived of the study, designed the experiments, and co-wrote the paper. A.W., T.C., D.P., A.H., and A.E.B. performed experiments and analyzed data. All authors reviewed the manuscript.

